# No evidence for extensive horizontal gene transfer in the genome of the tardigrade *Hypsibius dujardini*

**DOI:** 10.1101/033464

**Authors:** Georgios Koutsovoulos, Sujai Kumar, Dominik R. Laetsch, Lewis Stevens, Jennifer Daub, Claire Conlon, Habib Maroon, Fran Thomas, A. Aziz Aboobaker, Mark Blaxter

**Author notes:** corresponding author Mark Blaxter Ashworth Laboratories, The King’s Buildings, University of Edinburgh, Edinburgh EH9 3JT UK 01316506760.

## Abstract

Tardigrades are meiofaunal ecdysozoans that are key to understanding the origins of Arthropoda. Many species of Tardigrada can survive extreme conditions through cryptobiosis. In a recent paper (Boothby TC *et al* (2015) Evidence for extensive horizontal gene transfer from the draft genome of a tardigrade. *Proc Natl Acad Sci USA* 112:15976-15981) the authors concluded that the tardigrade *Hypsibius dujardini* had an unprecedented proportion (17%) of genes originating through functional horizontal gene transfer (fHGT), and speculated that fHGT was likely formative in the evolution of cryptobiosis. We independently sequenced the genome of *H. dujardini*. As expected from whole-organism DNA sampling, our raw data contained reads from non-target genomes. Filtering using metagenomics approaches generated a draft *H. dujardini* genome assembly of 135 Mb with superior assembly metrics to the previously published assembly. Additional microbial contamination likely remains. We found no support for extensive fHGT. Among 23,021 gene predictions we identified 0.2% strong candidates for fHGT from bacteria, and 0.2% strong candidates for fHGT from non-metazoan eukaryotes. Cross-comparison of assemblies showed that the overwhelming majority of HGT candidates in the Boothby *et al*. genome derived from contaminants. We conclude that fHGT into *H. dujardini* accounts for at most 1-2% of genes and that the proposal that one sixth of tardigrade genes originate from functional HGT events is an artefact of undetected contamination.

## Significance statement

Tardigrades, also known as moss piglets or water bears, are renowned for their ability to withstand extreme environmental challenges. A recently published analysis of the genome of the tardigrade *Hypsibius dujardini* (Boothby TC*, et al*. (2015) Evidence for extensive horizontal gene transfer from the draft genome of a tardigrade. *Proc Natl Acad Sci USA* 112:15976-15981), concluded that horizontal acquisition of genes from bacterial and other sources might be key to cryptobiosis in tardigrades. We independently sequenced the genome of *H. dujardini* and detected a low level of horizontal gene transfer. We show that the extensive horizontal transfer proposed by Boothby *et al*. was an artefact of a failure to eliminate contaminants from sequence data before assembly.

Tardigrades are a neglected phylum of endearing animals, also known as water bears or moss piglets (1). They are members of the superphylum Ecdysozoa (2), sisters to Onychophora and Arthropoda (3, 4). There are about 800 described species (1), though many more are likely to be as yet undescribed (5). All are small (tardigrades are usually classified in the meiofauna) and are found in sediments and on vegetation from the Antarctic to the Arctic, from mountain ranges to the deep sea, and in marine and fresh water environments. Their dispersal may be associated with the ability of many (but not all) species to enter cryptobiosis, losing almost all body water, and resisting extremes of temperature, pressure, and desiccation (6-9), deep space vacuum (10) and irradiation (11). Interest in tardigrades focuses on their utility as environmental and biogeographic markers, the insight their cryptobiotic mechanisms may yield for biotechnology and medicine, and exploration of their development compared to other Ecdysozoa, especially Nematoda and Arthropoda.

*Hypsibius dujardini* (Doyère, 1840) is a limnetic tardigrade that is an emerging model for evolutionary developmental biology (4, 12-21). It is easily cultured in the laboratory, is largely see-through (aiding analyses of development and anatomy; SI Appendix, Fig. S1), and has a rapid life cycle. *H. dujardini* is a parthenogen, with first division restitution of ploidy (22), and so is intractable for traditional genetic analysis, though reverse-genetic approaches are being developed (17). *H. dujardini* has become a genomic model system, revealing the pattern of ecdysozoan phylogeny (3, 4) and the evolution of small RNA pathways (23). *H. dujardini* is poorly cryptobiotic (24), but serves as a useful comparator for good cryptobiotic species (9).

Animal genomes can accrete horizontally transferred DNA, especially from germline-transmitted symbionts (25), but the majority of transfers are non-functional and subsequently evolve neutrally (dead-on-arrival, or doaHGT) (25-27). Functional horizontal gene transfer (fHGT) can bring to a recipient genome new biochemical capacities, and contrasts with gradualist evolution of endogenous genes to new function. The bdelloid rotifers *Adineta vaga* (28) and *Adineta ricciae* (29) have high levels of fHGT (˜8%), and this has been associated with both their survival as phylogenetically ancient asexuals and their ability to undergo cryptobiosis (28-32). Different kinds of evidence are required to support claims of doaHGT compared to fHGT. Both are supported by phylogenetic proof of foreignness, linkage to known host-genome-resident genes, *in situ* proof of presence on nuclear chromosomes (33), Mendelian inheritance (34), and phylogenetic perdurance (presence in all, or many individuals of a species, and presence in related taxa). Functional integration of a foreign gene into an animal genome requires adaptation to the new transcriptional environment including acquisition of spliceosomal introns, acclimatisation to host base composition and codon usage bias, and evidence of active transcription (for example in mRNA sequencing data) (35, 36).

Another source of foreign sequence in genome assemblies is contamination, which is easy to generate and difficult to separate. Genomic sequencing of small target organisms requires the pooling of many individuals, and thus also of their associated microbiota, including gut, adherent and infectious organisms. Contaminants negatively affect assembly in a number of ways (37), and generate scaffolds that compromise downstream analyses. Cleaned datasets result in better assemblies (38, 39), but care must be taken not to accidentally eliminate true HGT fragments.

A recent study based on *de novo* genome sequencing of *H. dujardini* came to the startling conclusion that 17% of this species’ genes arose by fHGT from non-metazoan taxa (13). Surveys of published genomes have revealed many cases of HGT (40), but the degree of fHGT claimed for *H. dujardini* would challenge accepted notions of the phylogenetic independence of animal genomes, and general assumptions that animal evolution is a tree-like process. The reported *H. dujardini* fHGT gene set included functions associated with stress resistance and a link to cryptobiosis was proposed (13). Given the potential challenge to accepted notions of the integrity and phylogenetic independence of animal genomes, this claim (13) requires strong experimental support. Here we present analyses of the evidence presented, including comparison to an independently generated assembly from the same *H. dujardini* strain, using approaches designed for low-complexity metagenomic and meiofaunal genome projects (38, 39). We found no evidence for extensive functional horizontal gene transfer into the genome of *H. dujardini*.

## Results and Discussion

**Assembly of the genome of *H. dujardini***. Using propidium iodide flow cytometry, we estimated the genome of *H. dujardini* to be ˜110 Mb, similar to a previous estimate (20). We sequenced and assembled the genome of *H. dujardini* using Illumina short-read technology. Detailed methods are given in *Supporting Information*. Despite careful cleaning before extraction, genomic DNA samples of *H. dujardini* were contaminated with other taxa. Adult *H. dujardini* have only ˜10^3^ cells, and thus a very small mass of bacteria would yield equivalent representation in raw sequence data. A preliminary assembly (called nHd.1.0) generated for the purpose of contamination estimation spanned 185.8 Mb. We expected assembly components deriving from the *H. dujardini* genome to have similar GC%, and to have the same coverage in the raw data (because each segment is represented equally in every cell of the organism). Contaminants may have different average GC%, and need not have the same coverage as true nuclear genome components. Taxon-annotated GC-coverage plots (TAGC plots or blobplots (38, 39)) were used to visualise the genome assembly and permitted identification of at least five distinct blobs of likely contaminant data with GC% and coverage distinct from the majority tardigrade sequence (Fig. 1A). Read pairs contributing to contigs with bacterial identification, and no mitigating evidence of tardigrade-like properties (GC%, read coverage and association with eukaryote-like sequences) were conservatively removed. There was minimal contamination with *C. reinhardtii*, the food source (41). Further rounds of assembly and blobplot analyses identified additional contaminant data (39) which was also removed. An optimised assembly, nHd.2.3, was made from the cleaned read set. Contigs and scaffolds below 500 bp were removed. Mapping of *H. dujardini* poly(A)+ mRNA-Seq (42) and transcriptome (12) data was equivalent between nHd.1.0 and nHd.2.3 (SI Appendix, Table S1), therefore we conclude that we had not over-cleaned the assembly.

**Figure 1.**
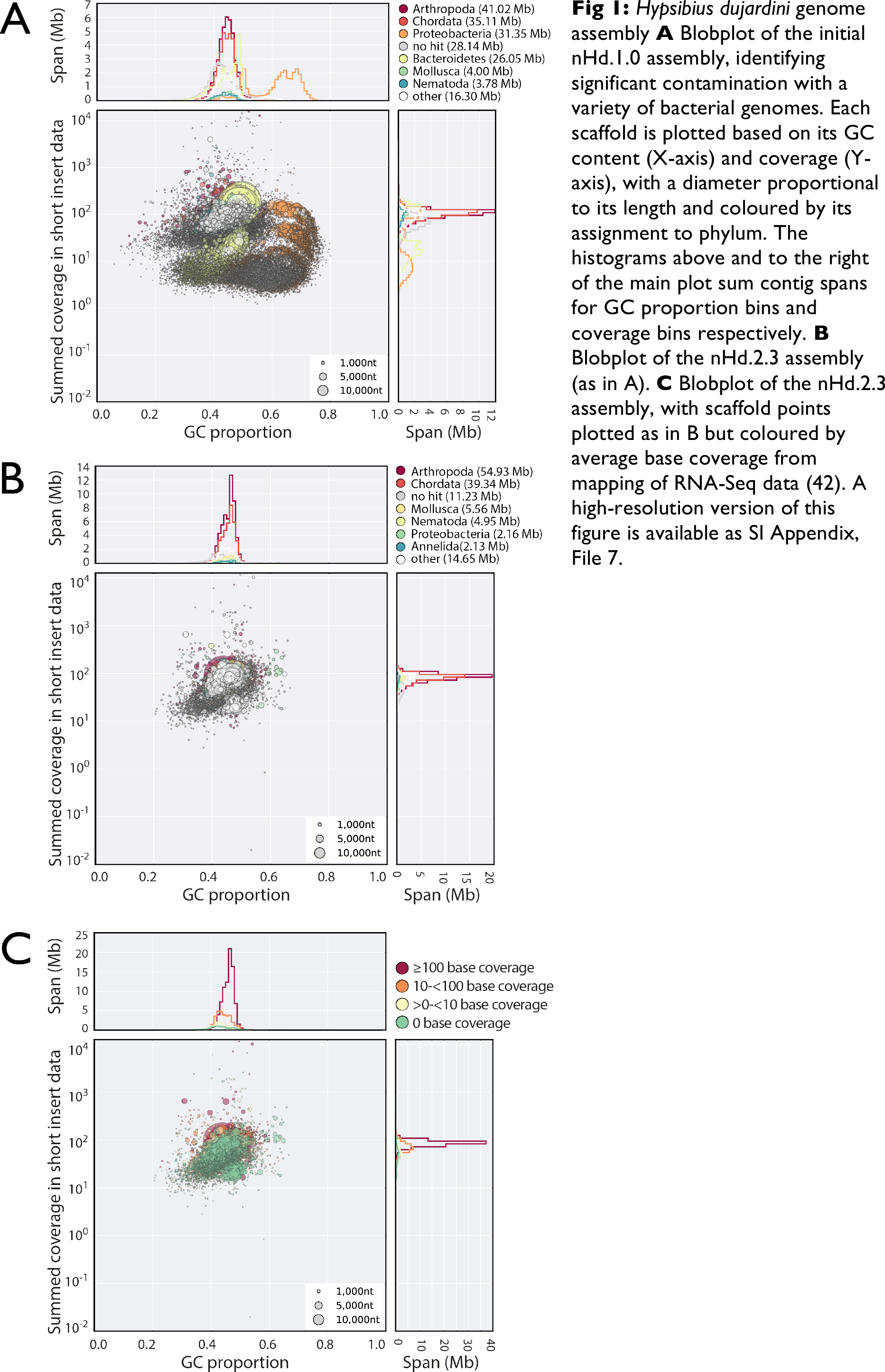
*Hypsibius dujardini* genome assembly. **A** Blobplot of the initial nHd.1.0 assembly, identifying significant contamination with a variety of bacterial genomes. Each scaffold is plotted based on its GC content (X-axis) and coverage (Y-axis), with a diameter proportional to its length and coloured by its assignment to phylum. The histograms above and to the right of the main plot sum contig spans for GC proportion bins and coverage bins respectively. **B** Blobplot of the nHd.2.3 assembly (as in A). **C** Blobplot of the nHd.2.3 assembly, with scaffold points plotted as in B but coloured by average base coverage from mapping of RNA-Seq data (42). A high-resolution version of this figure is available as SI Appendix, File 7.

The nHd.2.3 assembly had a span of 135 Mb, with an N50 length of 50.5 kb (Table 1). The assembly was judged relatively complete. It had good representation of a set of highly conserved, single-copy eukaryotic genes from the Core Eukaryotic Genes Mapping Approach (CEGMA) set (43), and these had a low duplication rate (1.3–1.5). A high proportion of *H. dujardini* mRNA-Seq (Fig. 1C) (42), transcriptome assembly (12), expressed sequence tags (ESTs), and genome survey sequences (GSSs) mapped to the assembly. We predicted a high-confidence set of 23,021 protein-coding genes using AUGUSTUS (44). The number of genes may be inflated because of fragmentation of the assembly, as 2,651 proteins lacked an initiation methionine, likely because they were at the ends of scaffolds, and were themselves short.

**Table 1.**
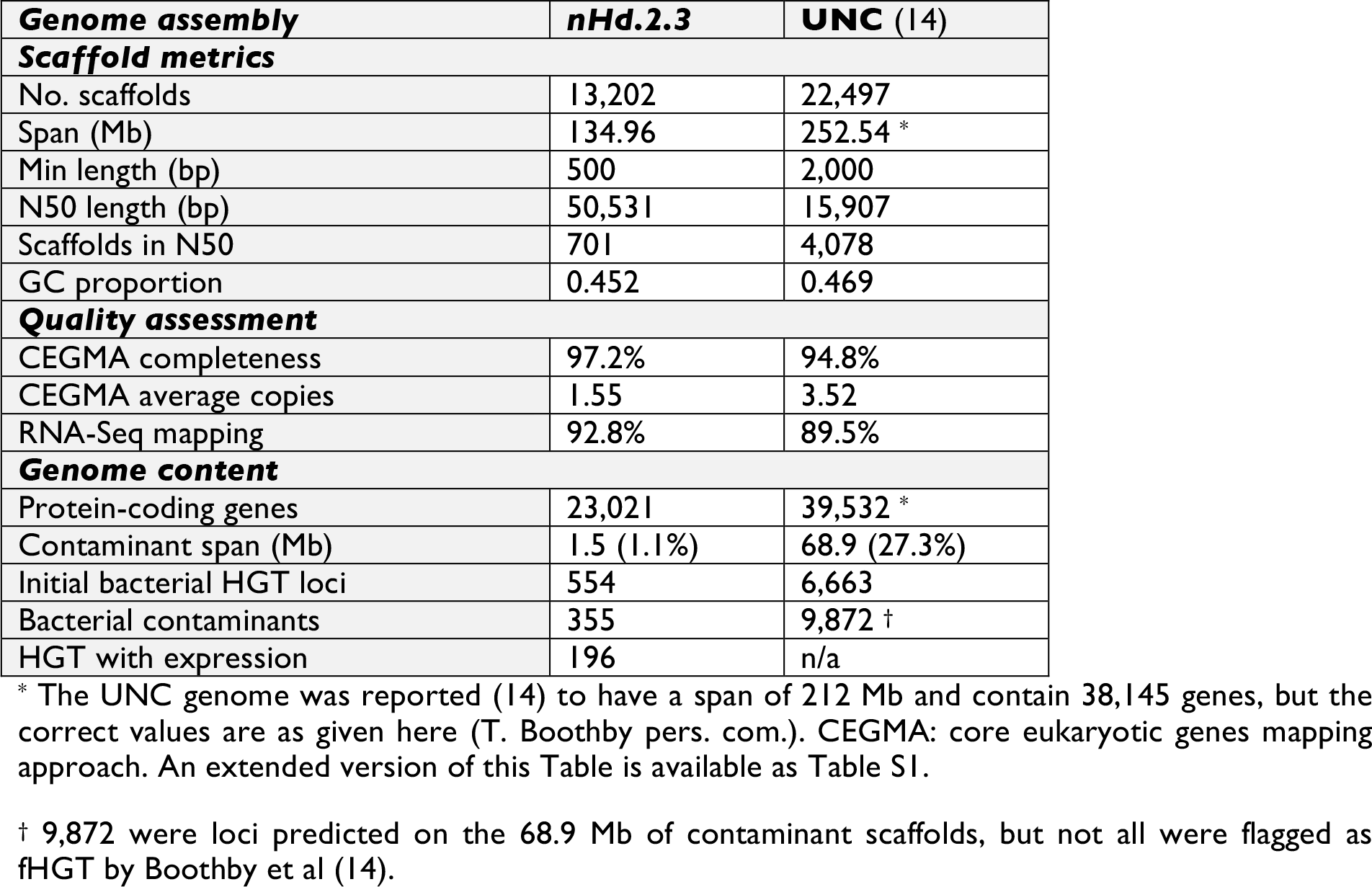
*Hypsibius dujardini* assembly comparison

Assembly of the *H. dujardini* genome was not a simple task, and the nHd.2.3 assembly is likely to still contain contamination. We identified 327 scaffolds (5.0 Mb) that had read coverage similar to *bona fide* tardigrade scaffolds but similarity matches to bacterial genomes (SI Appendix, File 2). Some of these scaffolds also encoded eukaryote-like genes, and may represent HGT or misassemblies. Some scaffolds (195 spanning 1.4 Mb) had only bacterial or no genes and were very likely to be contamination. We identified no scaffolds with matches to bacterial ribosomal RNAs (rRNAs) but did find an 11 kb scaffold with best matches to rRNAs from bodonid kinetoplastid protozoa. Two additional small scaffolds (6 kb and 1 kb) encoded kinetoplastid genes (a retrotransposon and histone H2A, respectively). No other genes were found on these scaffolds, and their high coverage likely resulted from the loci being multicopy in the source genome. The genome was made openly available to browse and download on a BADGER (45) server at http://www.tardigrades.org in April 2014 (SI Appendix, Fig. S3).

**Claims of extensive functional horizontal gene transfer into *H. dujardini*.** Boothby *et al*. (13) published an estimate of the genome of *H. dujardini*, referred to as the UNC [University of North Carolina] assembly hereafter, based on a subculture of the same culture sampled for nHd.2.3. They suggested that the *H. dujardini* genome was 252 Mb in span, that the tardigrade had 39,532 protein coding genes, and that over 17% of these genes (6,663) had been derived from extensive functional horizontal gene transfer (fHGT) from a range of prokaryotic and microbial eukaryotic sources. Given this claim, and the striking difference between the UNC assembly and our assembly, we set out to test the hypothesis that these “HGTs” were in fact unrecognised contamination in the UNC assembly.

Surprisingly, the UNC assembly had poorer metrics than nHd.2.3 (31) (Table 1), despite the application of two independent long read technologies (Pacific Biosciences [PacBio] and Moleculo) and equivalent short read data. Scaffold N50 length was one third that of nHd.2.3, despite UNC having discarded all scaffolds shorter than 2 kb. The UNC assembly span was 1.9 times that of nHd.2.3, in conflict with the UNC authors’ own (20) and our genome size estimates. The UNC protein prediction set was 1.7 times as large as that from nHd.2.3. UNC had good representation of CEGMA genes (Table 1), but contained over 3 copies on average of each single-copy locus. Such multiplicity of representation of CEGMA single-copy genes can arise (as the CEGMA gene set is not explicitly designed to exclude loci with bacterial homologues; see SI Appendix).

About one third of the span of UNC does not appear to be derived from the tardigrade. Many scaffolds had low coverage compared to *bona fide* tardigrade scaffolds (Fig. 2A), had different relative coverages in different libraries (SI Appendix, Fig. S4 B-D), were not represented in our raw data (Fig. 2B), and had overwhelmingly non-eukaryote taxonomic assignments (SI Appendix, File 3). The absence of all but marginal similarity to metazoan sequence also suggests that these contigs are not chimaeric co-assemblies. All the longest scaffolds in UNC were bacterial (Fig. 3A), and few bacterial scaffolds had read coverage support in both UNC and Edinburgh raw data (Fig. 3B). We identified 15 scaffolds in UNC with high-identity matches to rRNA genes from Armatimonadetes, Bacteroidetes, Chloroflexi, Planctomycetes, Proteobacteria and Verrucomicrobia (SI Appendix, Table S2). We also identified contamination that is likely to derive from other genomes. Two very similar UNC scaffolds (scaffold2445 and scaffold2691) both contained two, tandemly repeated copies of the rRNAs of a bdelloid rotifer related to *Adineta vaga*. We found a large number of additional matches to the *A. vaga* genome (28) in UNC, but these may be bacterial contaminants matching *A. vaga* bacterially-derived fHGT genes (28, 30). A total of 0.5 Mb of scaffolds had best sum matches to Rotifera rather than to any bacterial source (SI Appendix, File 3). Six mimiviral-like proteins were identified, five of which involved homologues of the same protein family (with domain of unknown function DUF2828). Mimiviruses are well known for their acquisition of foreign genes (46), and these scaffolds may derive from a mimivirus rather than the tardigrade genome. Overall, very few of the total fHGT candidates proposed by Boothby *et al*. (13) were in scaffolds that were not obviously contaminant (SI Appendix, File 4).

**Figure 2.**
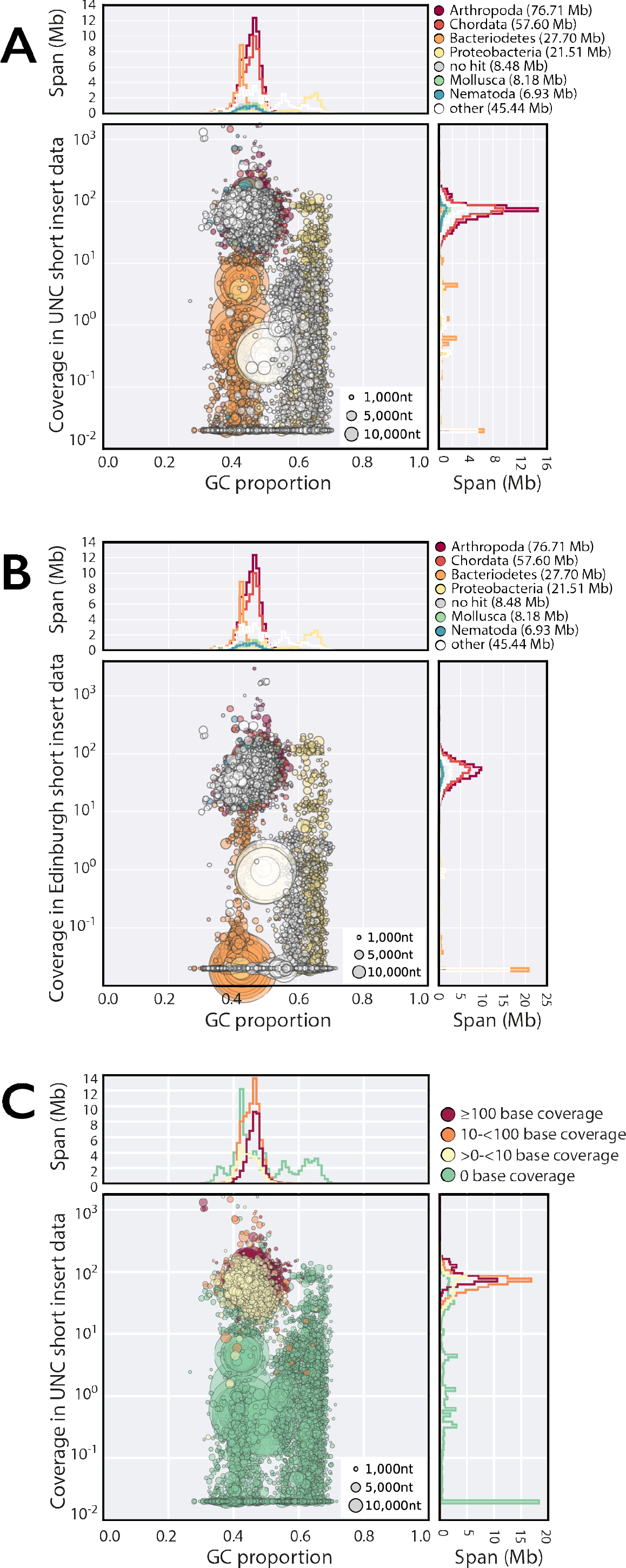
Contaminants in the UNC assembly. **A** Blobplot of the UNC assembly with coverage derived from pooled UNC raw genomic data. **B** Blobplot showing the UNC assembly with coverage derived from the Edinburgh short insert genomic data. **C** Blobplot (as in A) with the scaffold points coloured by average RNA-Seq base coverage. A high-resolution version of this figure is available as SI Appendix, File 7.

Presence of fHGT candidate transcripts in poly(A)-selected RNA is strong evidence for eukaryotic-like expression and integration into a host genome. We mapped *H. dujardini* mRNA-Seq data (42) to UNC (Fig. 2C). Only nine of the UNC scaffolds that had low or no read coverage in our raw genome data had appreciable levels of mRNA-Seq reads mapped (between 0.19 and 31 transcripts per million [tpm]). One of these (scaffold1161) contained two genes for which expression was >0.1 tpm, but all genes on this scaffold had best matches to Bacteria. The mRNA-Seq data thus gave no support for eukaryote-like gene expression from the low coverage, bacterial contigs in UNC.

Boothby *et al*. (13) assessed foreignness using an HGT index (47), and by analysing phylogenies of candidate genes. However these tests are only valid when there is independent evidence of incorporation into a host genome. Boothby *et al*. (13) assessed genomic integration of 107 candidate loci directly, using PCR amplification of predicted junction fragments. Most were adjudged confirmed, but no sequencing to confirm the expected amplicon sequence was reported. The 107 candidates were reported (13) to include 38 bacterial-bacterial and 8 archaeal-bacterial junctions (SI Appendix, Table S3). Our analyses identified 49 bacterial-bacterial junctions in their set, but these do not confirm HGT, as similar linkage would also be found in bacterial genomes. We found no expression of the 49 bacterial-bacterial junction loci, supporting assignment as contaminants rather than examples of fHGT.

Of the remaining 58 loci, only 51 were likely to be informative for HGT (SI Appendix, Table S3), as 7 were themselves or were paired with loci of unassigned taxonomic affinity. The informative loci included 24 prokaryotic to eukaryotic, 21 non-metazoan eukaryotic-metazoan and 6 viral-eukaryotic junction pairs. The UNC PacBio data confirmed only 25 of these junctions. All 58 loci had read coverage in Edinburgh raw data, and the same genomic environment was observed in nHd.2.3 for 51 loci (43 of which were HGT-informative). We found evidence of expression from 49 of these loci.

The UNC *H. dujardini* genome is thus poorly assembled and highly contaminated. Scaffolds identified as likely bacterial contaminants in UNC included 9,872 protein predictions (Table 1). Evidence for extensive fHGT is absent, and most candidates were not confirmed by PacBio data, our read data, or gene expression. We present a more detailed examination of each of Boothby *et al*’s claims for fHGT, including apparent congruence of codon usage and presence of introns, in SI Appendix.

**Low levels of functional horizontal gene transfer in *H. dujardini*.** We screened nHd.2.3 for loci potentially arising through HGT. As mapping of transcriptome data to nHd.2.3 was equivalent to the pre-cleaning nHd.1.0 assembly and better than the UNC assembly (SI Appendix, Table S1), the assembly has not been over-cleaned. Forty-eight nHd.2.3 scaffolds (spanning 0.23 Mb and including 41 protein-coding genes), had minimal coverage in UNC data (Fig. 3C), suggesting that these were contaminants. The remaining 13,154 scaffolds spanned 134.7 Mb. Of the 23,021 protein coding genes predicted, only 13,500 had sequence similarity matches to other organisms, and of these 10,161 had unequivocal signatures of being metazoan, with best matches to phyla including Arthropoda, Nematoda, Mollusca, Annelida, Chordata and Cnidaria. *A priori* these might be considered candidates for metazoan–metazoan fHGT. However, as *H. dujardini* is the first tardigrade sequenced, this pattern may just reflect the lack of sequence from close relatives. The remainder had best matches in a wide range of non-metazoan eukaryotes, frequently with metazoan matches with similar scores, and, for a few, bacterial matches. Some non-metazoan eukaryote-like proteins (e.g. the two bodonid-like proteins discussed above) may have derived from remaining contamination.

We found 571 bacterial-metazoan HGT candidates in nHd.2.3, of which 355 were on 166 scaffolds that contained only other bacterial genes. While some of these scaffolds also contained genes that had equivocal similarities, we regard them as likely remaining contaminants. Expression of these genes was in general very low (Fig. 3D) and we propose that these are “soft” candidates for fHGT. The remaining 216 HGT candidates were linked to genes with eukaryotic or metazoan classification on 162 scaffolds that had GC% and coverage similar to the tardigrade genome in both datasets (Table 2; SI Appendix, File 6). Most of these (196, 0.9% of all genes) had expression >0.1 tpm (Fig. 3D) and are an upper bound of “hard” candidates for fHGT. However, phylogenetic analyses identified only 55 (0.2% of all genes) with bacterial affinities (having only bacterial and no metazoan homologues, or where analysis of alignments including the closest metazoan homologues confirmed bacterial affinities; SI Appendix, File 6). We identified 385 loci (1.7% of all genes) most similar to homologues from non-metazoan eukaryotes (SI Appendix, File 6). Most (369) of these had expression >0.1 tpm, but phylogenetic analysis affirmed likely non-metazoan origin of only 49 of these (0.2% of all genes; SI Appendix, File 6).

**Table 2.**
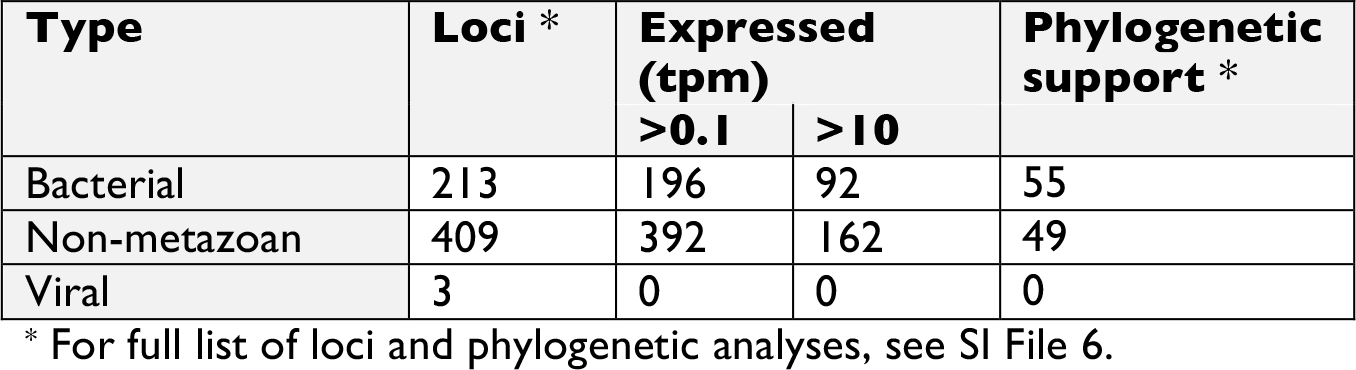
Putative HGT loci in *H. dujardini* nHd2.3

Within the high-coverage blob of assembly scaffolds supported by both Edinburgh and UNC raw data, blobtools analyses assigned 327 scaffolds as bacterial (black points in Fig. 3C). Fifty-two of these scaffolds were short (spanning 60.5 kb in total) and contained no predicted protein-coding genes, and 77 contained only predictions that were classified as eukaryote or unassigned. They were initially assigned as bacterial based on marginal nucleotide similarities to bacterial sequences. Many of the remaining 198 scaffolds were flagged in the gene-based analyses as containing fHGT candidates. Our assembly thus still contained contaminating sequences, mainly from bacteria but also including some from non-metazoan eukaryotes. *De novo* joint assembly of the Edinburgh and UNC datasets in the future will permit robust elimination of such “difficult” contamination, as well as definition of the correct genome span, true gene content and the contribution of HGT in *H. dujardini*.

**Figure 3.**
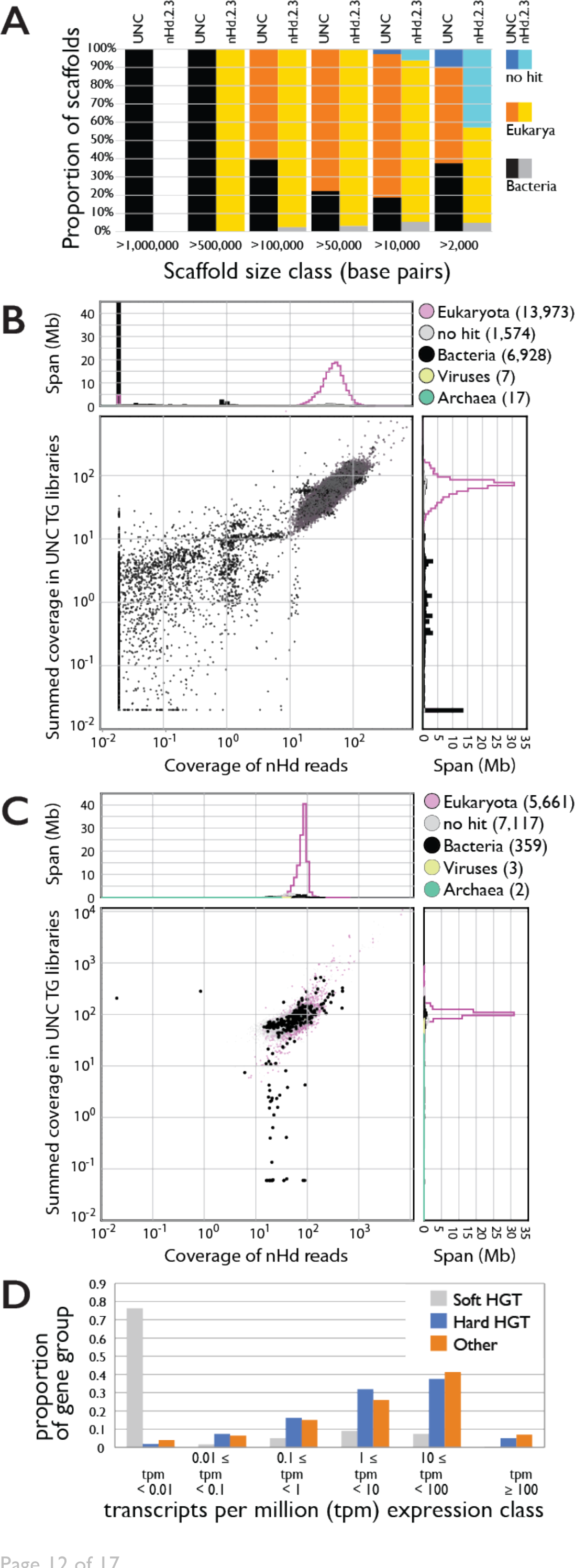
Identifying HGT candidates. **A** Stacked histogram showing scaffolds assigned to different kingdoms (Bacteria, Eukaryota, and “no hits”) in different length classes for UNC and nHd.2.3 assemblies. The nHd.2.3 assembly had no scaffolds >1 Mb, and all the longest scaffolds (>0.5 Mb) in the UNC assembly were bacterial. **B** Coverage-coverage plot of the UNC assembly using the Edinburgh short insert data (X-axis) and in the pooled UNC short insert data (Y-axis). **C** Coverage-coverage plot of the nHd.2.3 assembly as in B. **D** Expression of soft and hard HGT candidates, and all other genes, in the nHd.2.3 assembly. A high-resolution version of this figure is available in SI Appendix, File 7.

**Conclusions** We have generated a good draft genome for the model tardigrade *H. dujardini*. We identified areas for improvement of our assembly, particularly removal of remaining contaminant-derived sequences. We approached the data as a low complexity metagenomic project, and this methodology is going to be ever more important as genomics are used on systems difficult to culture and isolate. The blobtools package (38, 39) and related toolkits such as Anvi’o (48) promise to ease the significant technical problem of separating target genomes from those of other species. An Anvi’o analysis of these two tardigrade genomes confirms our finding that ˜70 Mb of the UNC assembly (including >96% of the proposed 6,663 HGT genes) derives from bacterial contamination (49).

Analyses of gene content and the phylogenetic position of *H. dujardini* and by inference Tardigrada are at an early stage, but are already yielding useful insights. Early, open release of the data has been key. The *H. dujardini* ESTs have been used for deep phylogeny analyses that place Tardigrada in Panarthropoda (3, 4), identification of a P2X receptor with an intriguing mix of electrophysiological properties (16), and for exploration of cryptobiosis in other tardigrade species (7, 8). The nHd.2.3 assembly was used for identification of opsin loci in *H. dujardini* (12).

Our assembly of the *H. dujardini* genome conflicts with the published UNC draft genome (13) despite being from the same original stock culture of *H. dujardini*. Our assembly had superior assembly and biological quality statistics but was ˜120 Mb shorter than UNC. About 70 Mb of the UNC assembly most likely derived from the genomes of several bacterial contaminants. The disparity between the non-contaminant span of the UNC assembly (˜180 Mb), our estimate of the genome (˜130 Mb) and direct densitometry estimates (80-110 Mb) may result from the presence of uncollapsed haploid segments. Resolution of this issue awaits careful reassembly.

We predicted a hugely reduced impact of predicted functional HGT: 0.2% to 0.9% of genes from nHd.2.3 had signatures of fHGT from bacteria, a relatively unsurprising figure. fHGT from non-metazoan eukaryotes into *H. dujardini* was less easily validated, but likely comprised a maximum of 0.2%. In *Caenorhabditis elegans*, *Drosophila melanogaster* and primates, validated bacterial fHGT loci comprise 0.8%, 0.3% and 0.5% of genes respectively (40). These mature estimates, from well-assembled genomes, are reduced compared to early guesses, such as the proposal that 1% of human genes originated through fHGT (50, 51). mRNA-Seq mapping shows that filtering did not compromise the assembly by eliminating *bona fide* tardigrade sequence. While some UNC fHGT candidates were confirmed, our analyses show that the UNC assembly is heavily compromised by sequences that derive from bacterial and other contaminants, and that the vast majority of the proposed fHGT candidates are artefactual.

## Experimental procedures

**Genome assembly and comparison to UNC assembly of *H. dujardini*.** The *H. dujardini* nHd.2.3 genome was assembled from Illumina short-insert and mate-pair data. We compared our assembly and that of Boothby *et al*. (13) by mapping raw read data and exploring patterns of coverage and GC% in blobtools (https://drl.github.io/blobtools) (38, 39) and exploring sequence similarity with BLAST and diamond. Details can be found in *Supporting Information*.

**Availability of Supporting Data.** Raw sequence read data have been deposited in SRA, dbGSS, and dbEST (SI Appendix, Table S4). Edinburgh genome assemblies have not been deposited in ENA, as we have no wish to contaminate the public databases with foreign genes mistakenly labelled as “tardigrade”. Assemblies (including GFF files, and transcript and protein predictions) are available at http://www.tardigrades.org and http://dx.doi.org/10.5281/zenodo.45436. Code used in the analyses is available from https://github.com/drl/tardigrade and https://github.com/sujaikumar/tardigrade.

**Competing interests.** The authors declare no competing interests.

**Author contributions.** AA, FT, JD, CC, HM and MB developed the *H. dujardini* culture system and prepared nucleic acids. EST and GSS sequencing was performed by JD, FT, CC, and HM. Genome assemblies were made by MB, SK, and GK. Analyses of the genomes were performed by MB, GK, SK, DRL and AA. LS constructed the BADGER instance and managed data. The manuscript was written by MB with input from all authors.

## Acknowledgments

The Edinburgh tardigrade project was funded by the BBSRC UK (grant reference 15/COD17089). GK was funded by a BBSRC PhD studentship. DRL is funded by a James Hutton Institute/School of Biological Sciences University of Edinburgh studentship. SK was funded by an International studentship and is currently funded by BBSRC award BB/K020161/1. LS is funded by a Baillie Gifford Studentship, University of Edinburgh. We thank Bob McNuff of Sciento for his inspired culturing of *H. dujardini*. Thanks to both reviewers who proposed changes that made this manuscript clearer. We especially thank a wide community of colleagues on twitter, blogs, and email for discussion of the results presented here (which were posted on bioRxiv for discussion: http://dx.doi.org/10.1101/033464) in the weeks since the publication of the UNC genome.

